# Functional and metabolomic analyses of brown adipose tissue during cold-deacclimation reveal rapid adaptations in N-acetylated amino acid metabolism

**DOI:** 10.1101/2025.09.23.678072

**Authors:** Chantal A. Pileggi, Ella McIlroy, Lauren M.K. Hamilton, Nidhi Kuksal, Luke S. Kennedy, Valeria Vasilyeva, Michel Kanaan, Ziyad El Hankouri, Yan Burelle, Miroslava Cuperlovic-Culf, Mary-Ellen Harper

## Abstract

Non-shivering thermogenesis (NST) in brown adipose tissue (BAT) is rapidly activated in cold environments and is an important thermoregulatory process. Despite the consensus that BAT is inactive under warm ambient temperatures, few studies have sought to examine the metabolic remodelling that occurs when recovering from the cold and re-acclimating to thermoneutral environments (28-32°C). To elucidate mitochondrial functional and structural aspects involved in BAT metabolic remodelling during cold deacclimation, we acclimated C57BL/6J mice to the cold (4°C) for 7 days, and subsequently transferred them to thermoneutrality (30°C) for 3 h, 12 h, 24 h, or 48 h. Comprehensive metabolic phenotyping analyses demonstrated elevated metabolic rates and high food intake during the cold acclimation period, which immediately decreased by ∼40% upon returning to thermoneutrality. High-resolution respirometry of saponin-permeabilized BAT revealed decreases in mitochondrial leak uncoupling by 24 h of cold deacclimation, which corresponded with gradual declines in mitochondrial protein content and UCP1 gene expression. Decreases in BAT mitochondrial content paralleled declines in protein content, as indicated by decreases in the mtDNA/nDNA ratio and mitochondrial surface area by 48 h of cold deacclimation. Metabolomic analysis of BAT from cold-acclimated mice and from mice deacclimated for 48 h at thermoneutrality revealed major changes in pathways related to amino acid metabolism, the tricarboxylic acid cycle (TCA), glutathione, and purine metabolism. Marked decreases in the abundance of N-acetylated amino acids in cold deacclimated mice corresponded with increased aminoacylase 1 (*Acy1*) expression. Together, these findings highlight the profound metabolic remodelling in BAT during thermogenesis and deactivation.

## Introduction

Brown adipose tissue (BAT) serves as an important thermoregulatory organ in homeothermic mammals, functioning to release energy as heat in cold environments through the process of non-shivering thermogenesis (NST) [1–3]. BAT is highly innervated and vascularized and is abundant in mitochondria that express high levels of the mitochondrial inner membrane protein, uncoupling protein 1 (UCP1) [4,5]. UCP1 mediates proton conductance across the inner mitochondrial membrane bypassing ATP synthase, thereby dissipating proton motive force and increasing substrate oxidation, which results in thermogenesis [6,7].

The molecular events involved in the initiation of cold-induced NST in small mammals are well-defined: Exposure to low ambient temperatures stimulates sensory thermal receptors in the skin to activate the sympathetic nervous system (SNS). The release of catecholamines, such as norepinephrine, stimulates β-adrenergic receptors on mature brown adipocytes [8,9]. Stimulation of β-adrenergic receptors increases cAMP concentrations and activates protein kinase A (PKA), which promotes lipolysis of triglycerides. Mobilized FFAs activate UCP1-mediated proton leak, leading to the increased uptake and rapid oxidation of fatty acids and glucose, which overall results in increased whole-body energy expenditure. Liberated fatty acids also stimulate PGC1α-mediated mitochondrial biogenesis and increase UCP1 expression [6,10,11]. The rise in cAMP also increases the conversion of thyroxine (T4) to 3,5,3′-triiodothyronine (T3) via the type 2 iodothyronine deiodinase (DIO2) selenoenzyme in BAT, and there is increased supply to BAT of thyroid hormone through the circulation via norepinephrine [12–14]. Within mature brown adipocytes, mobilized FFAs activate UCP1-mediated proton leak, leading to the increased uptake and rapid oxidation of fatty acids and glucose. In rodent studies, sustained or repeated bouts of cold exposure over periods of >5-7 days is termed cold acclimation. Cold acclimation results in an adapted metabolic state which includes increased vascularization and sympathetic innervation to support brown adipocyte proliferation and differentiation underlying BAT hyperplasia and hypertrophy [15–17]. Rapid increases in BAT metabolic activity precede the molecular remodelling of brown adipocytes during cold-acclimation, consistent with the idea that certain metabolites may function as signaling molecules that not only support acute thermogenic responses but also drive molecular remodeling of the tissue to support increased thermogenic capacity [18,19].

Despite the consensus that BAT is inactive under warm ambient environments, few studies have sought to examine the molecular events and metabolic remodelling that occur when recovering from the cold and re-acclimating to thermoneutral environments (28-32°C for mice) [20,21]. Understanding the profound metabolic remodelling that leads to the restitution of active thermogenic BAT into inactivated BAT is crucial for a comprehensive understanding of adaptive thermogenesis processes overall. Excessive or inappropriate cessation of BAT thermogenesis could potentially lead to hypothermia, for example, in neonates, who have much higher levels of BAT than adults. Elucidating the mechanisms underlying how BAT thermogenesis is inhibited may provide novel insights into ways through which low levels of BAT thermogenesis in certain metabolic diseases can be increased to prevent or treat these diseases. Here, we present a time-resolved analysis of the metabolic and structural remodelling that occurs during BAT inactivation in response to cold deacclimation. We hypothesized that cold deacclimation would result in remodelling that corresponds with rapid decreases in BAT mitochondrial content and oxidative capacity. Our results demonstrate that rapid changes in whole-body metabolism in response to cold deacclimation precede the molecular remodelling of BAT mitochondria and suggest that N-acetylated amino acids may function as metabolite reporter candidates to drive responses in BAT.

## Results

### Resting metabolic rate rapidly decreases during cold-deacclimation at thermoneutrality

To assess the metabolic adaptations of cold deacclimation, C57BL/6J mice were acclimated to the cold (4°C for 7 days) and then transferred to a thermoneutral climate (30°C) for 5 days (**Figure 1A)**. Consistent with previous findings [6,22], *ad libitum* food intake was high during cold acclimation and immediately decreased upon returning to thermoneutrality (**Figure 1B**). As anticipated, the decrease in food intake upon reacclimating to thermoneutrality coincided with a rapid ∼55% decline in metabolic rate (VO_2_, **Figure 1C**). Metabolic rates remained slightly elevated during the first 2 days of deacclimating at thermoneutrality and gradually declined by an additional ∼15% by day 4 of deacclimation (**Figure 1C**). Respiratory exchange ratio (RER, **Figure 1D**) gradually increased during the deacclimation period, indicating a relative shift from lipid to carbohydrate substrate oxidation. While total activity of mice in the chambers was not different in response to the acclimation protocol, ambulatory activity was low throughout cold exposure and increased during deacclimation. (**Figure 1E,F**) [23].

**Figure 1.**
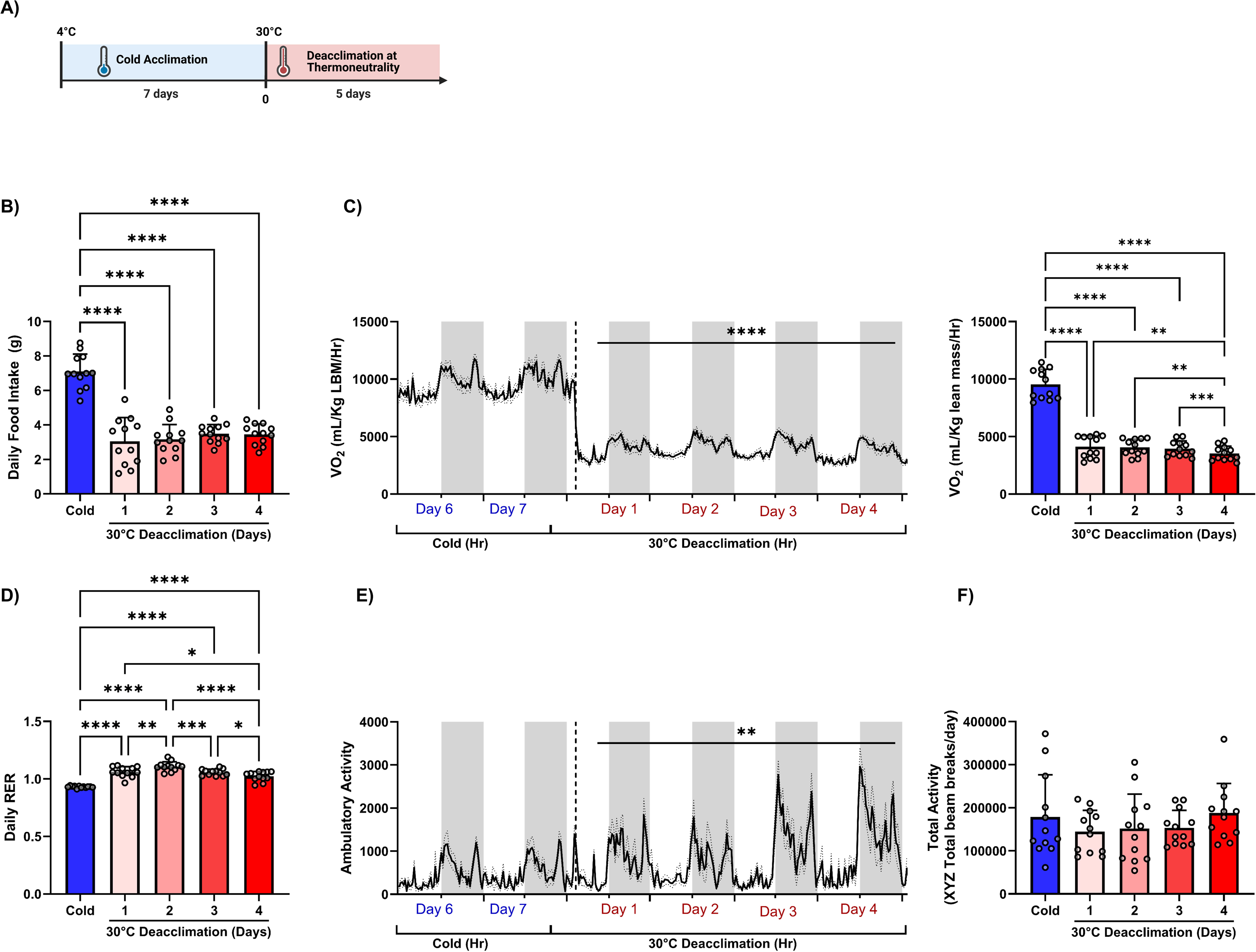
Resting metabolic rate rapidly decreases during cold-deacclimation. **(A-F)** Comprehensive metabolic phenotyping analyses was measured using a fully automated CLAMS system (Columbus Instruments) during cold acclimation (4°C) and subsequent deacclimation at thermoneutrality (30°C). **(A)** Schematic summary of the time points for sample collection. Diagram created with BioRender.com. **(B)** Daily *ad libitum* cumulative food intake. **(C)** Average hourly metabolic rates (VO2; normalized to lean body mass (LBM)). **(D)** Respiratory exchange ratio (RER; VCO2/VO2). **(E)** Ambulatory activity was calculated as the sum of ambulatory beam breaks in the x, y, and Z planes. **(F)** Total activity was calculated as the sum of all beam breaks in the x, y, and Z planes. Comparisons between timepoints were determined using a one-way RM ANOVA with Tukey post-hoc tests, *<p<0.05 **p<0.01, ***p<0.001, ****p<0.0001. All values are presented as means ± SD.

### Mitochondrial oxidative capacity and content regress in parallel during cold deacclimation

To elucidate mitochondrial functional and structural aspects involved in BAT metabolic remodelling, we acclimated C57BL/6J mice to the cold (4°C) with *ad libitum* access to food and water for 7 days and subsequently transferred them to thermoneutrality (30°C) for 3 h, 12 h, 24 h, or 48 h (**Figure 2A**). We selected these timepoints, as metabolic remodelling is initiated within the first 3 days of temperature acclimation [23]. Deacclimation at thermoneutrality did not impact body weights or the weights of interscapular BAT (iBAT) or inguinal white adipose tissue (**Figure 2B and S1A-C**).

**Figure 2.**
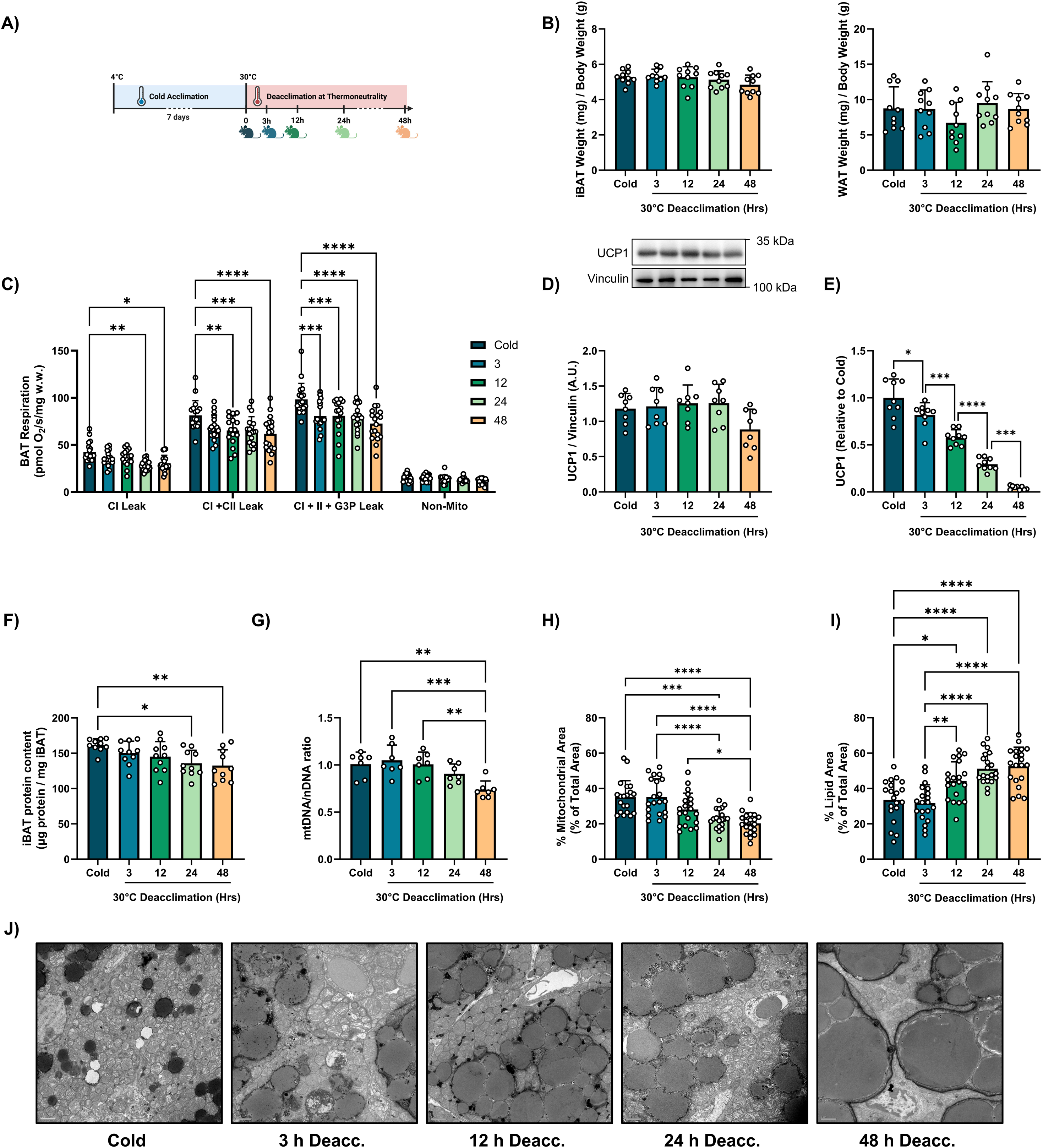
Capacity for mitochondrial uncoupling and mitochondrial content regress in parallel during cold deacclimation. **(A)** Schematic summary of the time points for sample collection. C57BL/6J mice were acclimated to the cold (4°C) for 7 days with *ad libitum* access to food and water and subsequently transferred to thermoneutrality (30°C) for 3 h, 12 h, 24 h, or 48 h. Diagram created with BioRender.com. **(B)** High-resolution respiratory flux per mg of saponin-permeabilized iBAT. Uncoupled respiration was measured in the presence of oligomycin and octanoylcarnitine. Complex I (CI) leak respiration was measured after the addition of malate-pyruvate-glutamate, and CI + CII leak and CI + II + G3P leak respiration were measured following the sequential additions of succinate and glycerol-3-phosphate for respectively, (n=16-18/group). **(C)** Immunoblotting of UCP1 protein expression (n=8/group). **(D)** Quantitative PCR (qPCR) of *Ucp1* gene expression, (n=9/group). **(E)** BAT protein concentration (per mg tissue) was measured using a BCA assay in protein lysates, (n=10/group). **(F)** qPCR was used to determine the mtDNA:nDNA ratio (mt-ND1:n-HK2) (n=7/group). **(G-J)** TEM images were analyzed by quantitative morphometry for (G) mitochondrial surface area and (H) lipid droplet surface area, (J) Representative TEM images, (n=5 TEM images from 4 mice/group). See also Figure S1. Comparisons between timepoints were determined using a one-way ANOVA with Tukey post-hoc tests, *<p<0.05 **p<0.01, ***p<0.001, ****p<0.0001. All values are presented as means ± SD.

We sought to determine the functional changes in iBAT oxidative capacity using high-resolution respirometry (HRR) conducted on saponin-permeabilized iBAT. To assess respiration attributable to mitochondrial proton leak uncoupling, HRR assays were conducted in the presence of oligomycin and octanoyl carnitine to support fatty acid-driven uncoupled respiration. Upon deacclimating from the cold at thermoneutrality, decreases CI + CII leak respiration were apparent by 12 h of deacclimation, and complex I (CI) leak respiration gradually declined by 24 h (**Figure 2C**). However, more rapid declines were observed in the leak respiration linked to the oxidation of glycerol-3-phosphate (G3P) by the FAD-linked mitochondrial glycerol-3-phosphate dehydrogenase (mGPDH), which were significant by 3 hours (**Figure 2C**). Importantly, the activity of mGPDH is relatively high in iBAT compared to other tissues, and mGPDH is transcriptionally controlled by triiodothyronine (T3) [24,25]. As the esterification/acylation of G3P is a major rate-controlling step for the synthesis of glycerophospholipids and triacylglycerols [26], the rapid decline in G3P-supported respiration may indicate a shift towards lipid synthesis rather than oxidation.

Despite the decreases in leak respiration, protein expression of UCP1 did not differ between time points of cold deacclimation (**Figure 2D**). In contrast, *Ucp1* mRNA expression was rapidly downregulated (**Figure 2E**). UCP1 protein has a half-life of ∼7 days *in vivo*, with differences paralleling changes in iBAT protein content [27]. Indeed, decreases in iBAT protein content were evident after 48 h of cold deacclimation (**Figure 2F).** In line with the rapid downregulation in mitochondrial respiration, markers of BAT activation, *Dio2* and ELOVL fatty acid elongase 3 (*Elvol3*) decreased by 3 h of cold deacclimation, whereas regulators of BAT activity PR domain containing 16 (*Prdm16*) and cell death inducing DFFA like effector a (*Cidea*) did not change across these selected time points of cold deacclimation (**Figure S1D**).

Changes in iBAT mitochondrial content paralleled the declines in protein content, as indicated by decreases in the mtDNA/nDNA ratio, citrate synthase activity, and mitochondrial surface area and number in electron micrographs by 48 h of cold deacclimation (**Figure 2G-2J, S1E,G**). The decreased mitochondrial content was partly due to the rapid downregulation of PGC1α-mediated mitochondrial biogenesis, as evidenced by decreases in *Pparg1ca* gene expression (**Figure S1F**). While gene expression of the nuclear respiratory factor 1 (*Nrf1*) did not differ in response to cold deacclimation, *Nrf2* was lower by 48 hours of cold deacclimation **(Figure S1F**). As mitochondrial content decreased during cold deacclimation, lipid droplet size gradually expanded (**Figure 2I, 2J, S1H,I**) likely owing to reduced lipolysis and triglyceride turnover [28].

### Cold deacclimation induces extensive remodelling of BAT metabolism, including decreases in glutathione and TCA cycle metabolism

Studies assessing the iBAT metabolome during cold exposure and acclimation have demonstrated the profound capacity of metabolic remodelling of iBAT. For example, there are rapid increases in the abundance and oxidation of fatty acids, amino acids, and glucose. To determine the impact of cold deacclimation, we assessed changes in the iBAT metabolome using quantitative ion-pairing liquid chromatography-mass spectroscopy (LCMS) in cold-acclimated mice and mice reacclimated at thermoneutrality for 48 h (**Figure 3A**). Principal-component analysis demonstrated clear separation of the clustered iBAT metabolite profiles from cold-acclimated mice and 48 h deacclimated mice (**Figure 3B**). After false discovery rate correction, we identified 53 out of 136 detected metabolites that differed between cold-acclimated and deacclimated mice (FDR<0.05) **(Figure 3C**). The machine learning feature selection method, ReliefF, identified N-acetyl amino acids (Glu, Met, Ser, Gln), guanosine, orotic acid, and N-carbamoyl-DL-aspartic acid as key metabolites responsible for discriminating the cold-acclimated vs. cold-deacclimated BAT metabolic phenotypes (**Figure 3D**). Metabolite set enrichment analysis (MSEA) demonstrated that cold deacclimation induced changes in pathways related to amino acid metabolism (aspartate, glycine/serine), glycolysis/gluconeogenesis, purine and nucleotide sugar metabolism, and the TCA cycle (**Figure 3E**).

**Figure 3:**
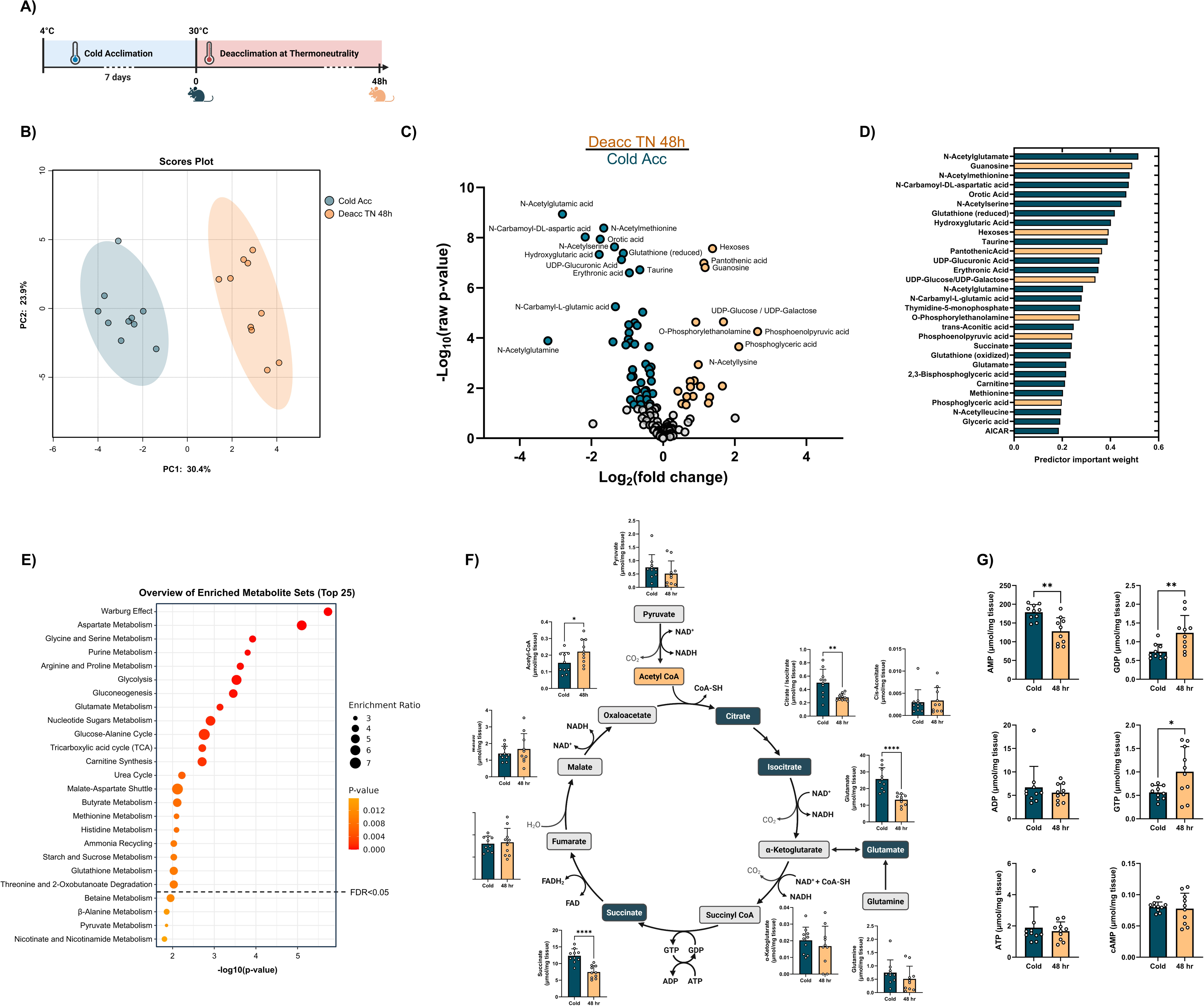
Metabolic profiling reveals cold deacclimation induces changes in the metabolism of amino acids, glutathione, and in TCA cycle activity. **(A)** Schematic summary of the time points for sample collection for metabolomic analyses of BAT from cold acclimated and 48h deacclimated mice, (n=10/group). Diagram created with <u>BioRender.com</u>. **(B)** Principle component analysis of independent samples demonstrating separation between the BAT metabolome profiles. **(C)** Volcano plots of BAT metabolite profiles from cold acclimated mice and mice deacclimated at thermoneutrality for 48h. Indicated in colour are metabolites with p<0.05 and log_2_ fold change over 1. Labelled are the top 20 metabolites. **(D)** Selection of the most significantly different features using the machine learning method ReliefF. **(E)** Metabolite set enrichment analyses of the metabolites altered between BAT from cold acclimated and deacclimated mice. **(F-G)** Quantitative analysis of individual metabolites relating to **(F)** the TCA cycle and **(G)** purine metabolism. Comparisons between groups were determined using Metaboanalyst. Comparisons for individual metabolites were determined using a two-tailed Student’s t-test. ∗p < 0.05, ∗∗p < 0.01, ∗∗∗p < 0.001, and ∗∗∗∗p < 0.0001.

Substrate metabolism during cold deacclimation appears to favour glycogenesis and gluconeogenesis during cold-deacclimation, as evident by higher abundances of UDP-glucose and hexoses available for glycolysis entry and a lower ratio of pyruvate/lactate (**Figure S2A,B**). Similarly, analyses of TCA metabolites revealed decreased concentrations of citrate and succinate during cold deacclimation, suggesting an overall decrease in TCA cycle activity during the deacclimation period (**Figure 3F**). Lower succinate in cold-deacclimated BAT is interesting given that accumulation of succinate can induce UCP1-mediated respiration in brown adipocytes, independent of adrenergic signalling [18]. Moreover, purine nucleotides ATP, ADP, GTP, and GDP can bind directly to UCP1 and inhibit proton leak uncoupling. Consistent with decreased UCP1 activity, the abundance of both GDP and GTP was higher in BAT from cold-deacclimated mice, whereas AMP decreased in abundance in cold deacclimated BAT (**Figure 3G**).

Other notable differences include increased abundance of pantothenic acid in cold-deacclimated BAT (**Figure S2C**), indicating increased availability of CoA. Pantothenic acid is required for adipocyte browning and the induction of UCP1 expression [29,30]. Pantothenate kinase 1 (PANK1), the rate-limiting enzyme for CoA synthesis, correlates with UCP1 expression in human BAT [31,32]. Glutathione and metabolites such as serine and methionine, which are involved in the synthesis of cysteine, the rate-limiting amino acid in glutathione synthesis, were lower in cold-deacclimated BAT (**Figure S2D**). ROS levels are high in BAT during UCP1-mediated mitochondrial uncoupling [33], and thus, the decreased abundance of GSH-related metabolites likely reflects the decreased need for the antioxidant properties of GSH during cold deacclimation. The abundance of carnitine decreased during cold deacclimation (**Figure S2E**), which, together with the decrease in citrate, potentially indicates a decrease in futile fatty acid cycling. Indeed, gene expression of the *de novo* lipogenesis enzyme ATP-citrate lyase (ACLY), which generates acetyl-CoA from citrate, decreased by 48 hours of cold deacclimation (**Figure S2F**) [34,35]. However, gene expression of acetyl-CoA synthetase 2 (ACSS2), which produces acetyl-CoA from acetate for lipid synthesis, did not change during the cold deacclimation period (**Figure S2G)**.

### Aminoacylase 1 may regulate the abundance of N-acetylated amino acids during cold acclimation and deacclimation

N-acetyl amino acids were the top metabolites altered during cold-deacclimation (**Figure 4A**). Tracing experiments using [13C]-glucose in iBAT of mice acclimated at cold, room, or thermoneutral temperatures have similarly identified N-acetylated amino acids as thermogenic reporter metabolite candidates, and the observed [13C]-glucose labelling at M+2 suggests that the acetyl group is derived from glucose [19]. Distance correlation analysis between N-acetyl amino acids and all other metabolites revealed that the abundances of N-acetyl amino acids at 48h of deacclimation strongly correlated with metabolites involved in the pentose phosphate pathways, gluconeogenesis (**Figure 4B,C**).

**Figure 4:**
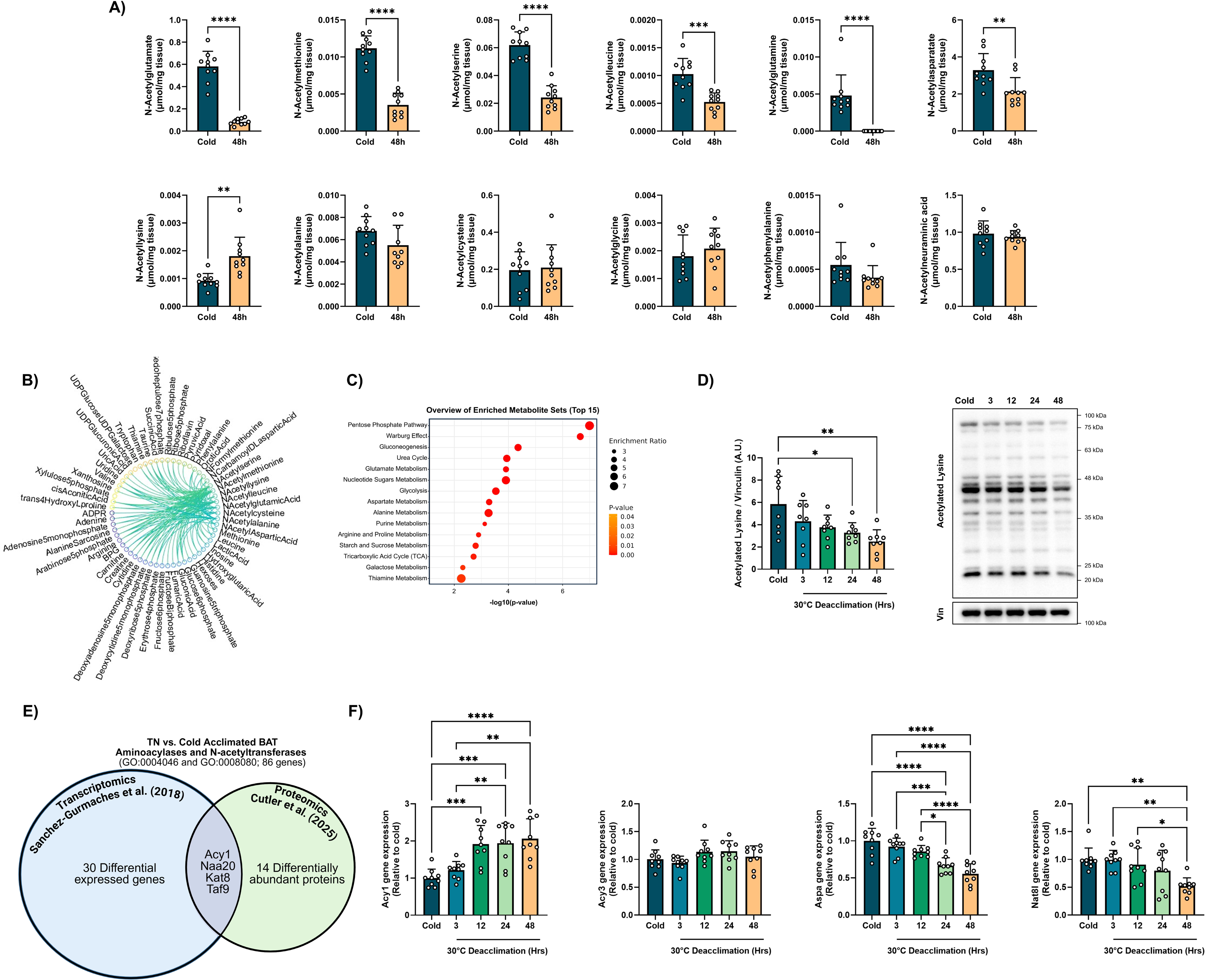
Aminoacylase 1 likely regulates the abundance of N-acetylated amino acids during cold acclimation and deacclimation. **(A)** Individual metabolite abundance of select N-acetylated amino acids. Comparisons were determined using a two-tailed Student’s t-test. **(B)** Distance correlations of the N-acetylated amino acids relating to the other measured metabolites, (n=10/group). Shown are metabolites that have a significant difference between iBAT from cold and 48 h deacclimated mice, where in 48 h deacclimated BAT they have correlation over 0.7 (p< 0.003) with any of the N-Acetylated-AAs and no such correlation in the cold BAT. **(C)** Metabolite set enrichment analyses of the metabolites that correlate with the N-acetylated amino acids at 48 h of cold deacclimation, but not in cold exposed BAT. **(D)** Immunoblotting of acetylated lysine residues on proteins, (n=8/group). Comparisons between timepoints were determined using a one-way ANOVA with Tukey post-hoc tests. **(E)** Bioinformatic analysis of published RNA-seq [37] and proteomics [38] datasets to identify proteins involved in the (de)acetylation of proteins and amino acids during cold acclimation using the GO terms N-acetyltransferase activity (GO:0008080) and aminoacylase activity (GO:0004046). Four proteins were shared between datasets: Aminoacylase 1 (Acy1), Nα-acetyltransferase 20 (Naa20), lysine acetyltransferase 8 (Kat8), and TATA-box binding protein associated factor 9 (Taf9). **(F)** Quantitative PCR (qPCR) of aminoacylase 1 (Acy1) and 3 (Acy3), aspartoacylase (Aspa), and N-Acetyltransferase 8-Like (Nat8l), (n=9/group). Comparisons between timepoints were determined using a one-way ANOVA with Tukey post-hoc tests. All values are presented as means ± SD. *<p<0.05 **p<0.01, ***p<0.001, ****p<0.0001.

High levels of N-acetylated amino acids during cold exposure may arise from the proteolytic degradation of N-terminal acetylated proteins or the transfer of an acetyl group to free amino acids via N-acetyltransferases. Thus, we sought to explore the mechanisms underlying the rapid change in abundance of N-acetyl amino acids. Interestingly, N-acetyllysine was the only N-acetylated amino acid to increase in abundance with cold-deacclimation (**Figure 4A**). This increase in N-acetyllysine corresponded with a decrease in protein lysine acetylation during cold deacclimation (**Figure 4D**), indicating an increase in protein degradation throughout the cold deacclimation period. Thus, we hypothesized that the decrease in N-acetylated amino acids during cold deacclimation was due to decreased activity of N-acetyltransferases, possibly combined with increased activity of zinc-dependent aminoacylases (ACY), which can deacetylate a wide range of Nα-acetylated amino acids, by hydrolyzing N-acetyl L-amino acids into free amino acids [36].

We then leveraged publicly available published RNA-seq [37] and proteomics [38] datasets to identify proteins involved in the (de)acetylation of proteins and amino acids during cold acclimation using the GO terms N-acetyltransferase activity (GO:0008080) and aminoacylase activity (GO:0004046). Of the 86 genes mapped to these terms, 30 genes were differentially expressed, and 14 proteins differed in abundance (**Figure 4E, Table S1**). Four proteins were shared between datasets: Aminoacylase 1 (Acy1) was lower in BAT from cold acclimated vs.TN mice, whereas the catalytic subunit of NatB complex, Nα-acetyltransferase 20 (Naa20), lysine acetyltransferase 8 (Kat8), and TATA-box binding protein associated factor 9 (Taf9) were higher in BAT from cold acclimated vs.TN **(Figure 4E, Table S1**).

We concentrated on aminoacylase expression, as ACY1 is strongly associated with the abundances of several N-acetyl amino acids in human plasma (85, 86). There are 3 family members, which differ in amino acid specificity: ACY1 deacetylates neutral aliphatic N-acyl and N-acetyl-α-amino acids and mercapturic acids; ACY2 or aspartoacylase (ASPA) specifically deacetylates N-acetyl-aspartate; and ACY3 preferentially deacetylates Nα-acetylaromatic amino acids. Gene expression of *Acy1* increased during cold deacclimation, whereas *Aspa* decreased during cold deacclimation (**Figure 4F**), whereas *Acy3* expression did not change during cold deacclimation. Gene expression of *Aspa* decreased during cold deacclimation, which corresponded with reduced expression of N-acetyltransferase 8-Like (*Nat8l*), which synthesizes N-acetylaspartate from acetyl-CoA and aspartate (**Figure 4F**). Taken together, these data indicate that aminoacylase expression may be sensitive to environmental temperature and may contribute to the marked decreases in N-acetylated metabolites.

## Discussion

Brown adipose tissue is characterized by significant morphological and functional plasticity in response to changes in ambient temperatures. Most studies have focused on processes occurring during cold acclimation, and relatively little is known about mechanisms elicited during deacclimation from the cold. Here, we conduct a time-resolved analysis of iBAT deactivation during the transition to thermoneutrality following 7 days of cold acclimation. Our results demonstrate that whole-body metabolism is rapidly decreased when cold-acclimated mice are returned to thermoneutrality, and analyses of iBAT structure and function revealed the gradual remodelling of BAT mitochondria. Analyses of the iBAT metabolome highlighted the marked alterations in amino acid metabolism, glutathione, and TCA cycle between cold acclimated and 48 h deacclimated mice. Striking decreases in N-acetylated amino acids in 48h deacclimated BAT indicate that select N-acetylated amino acids act as thermogenic reporters and are possibly regulated by temperature-induced alterations in aminoacylase expression.

The duration of the cold exposure affects the pace at which BAT is inactivated when animals return to warm environments, with longer periods of cold acclimation often resulting in slower regression of BAT, with full deacclimation occurring after 2-3 weeks [39,40]. Upon returning to a warm environment after a period of cold acclimation, decreases in SNS activity, blood flow, and substrate uptake induce the dynamic metabolic remodelling of BAT [41,42]. However, cold-induced metabolic adaptations may persist for several days after the cold exposure period has concluded, such as transient improvements in glucose homeostasis and redox homeostasis [43–48]. Despite immediate drops in whole-body energy expenditure, our data demonstrating a higher metabolic rate that persists for 2 days after returning to thermoneutrality further highlights the metabolic costs of adaptations from cold acclimation. Studies in humans have similarly observed that many of the cold-induced metabolic effects do not immediately subside fully during the initial re-warming period [49]. Specifically, sustained increases in glucose uptake, heat production, and substrate oxidation remain elevated for several hours to days after returning to warm environments [49,50]. The observed hyperphagic response in cold-acclimated mice is thought to be a result of substantially increased metabolic demand to maintain energy reserves for oxidation and to induce a degree of diet-induced thermogenesis via the thermic effect of feeding, which provide additional thermogenic effects [51–54]. Our finding that food intake decreases within the first 24 hours of returning to a thermoneutral environment is consistent with the important role of neuronal control of food intake to maintain energy homeostasis. Similarly, lower ambulatory and rearing activity in the cold may be attributable to behavioural and physiological responses to maintain body temperature by limiting or expanding body surface area exposed to ambient temperatures [55].

In contrast to the immediate declines in whole-body energy expenditure, permeabilized iBAT did not exhibit immediate decreases capacity for uncoupled mitochondrial respiration. Blood flow and glucose uptake in iBAT rapidly decrease in BAT within 2-6 hours (17, 18), suggesting that a decreased supply of substrates to BAT mitochondria may underlie the reductions in metabolic capacity. BAT oxygen consumption strongly correlates with blood flow to BAT [31,56,57]. The functional changes in mitochondrial oxidative capacity following the cessation of cold-induced thermogenesis appear to coincide with the gradual declines in mitochondrial content, UCP1 protein expression and BAT protein content, which is consistent with previous findings [27,39,48,58,59]. The declining BAT mitochondrial content observed during cold deacclimation is accompanied by increases in unilocular lipid droplet formation and glycogen repletion [39,47,60,61], consistent with decreased demand for substrate oxidation. Together, these results suggest that a decreased supply of substrates for oxidation, for example due to decreased supply from the circulation or to decreased SNS stimulation of cellular lipolysis, may drive immediate temperature-induced decreases in BAT oxidative capacity and metabolic remodelling.

Sophisticated *in vivo* metabolite tracing experiments have demonstrated that glucose and fatty acids primarily fuel BAT during thermogenesis [62–64]. Cytosolic fatty acid synthesis also increases during thermogenesis, resulting in the futile cycling of fatty acids [65]. Citrate generated by the TCA cycle is exported to the cytosol via the citrate-malate shuttle and subsequently metabolized by ATP-citrate lyase (ACLY) to synthesize acetyl-CoA for *de novo* lipogenesis via acetyl-CoA carboxylase (ACC1) and fatty acid synthase (FASN) [35,61,66]. The resulting acyl-carnitines can then return to the mitochondria for oxidation [35]. Our observations of decreased citrate and carnitine abundance and decreased gene expression of ATP-citrate lyase by 48 h of deacclimation are in line with the notion that the futile cycling between fatty acid synthesis and oxidation is reduced following the cessation of the cold stimulus. The rapid changes in BAT metabolic capacity precede the molecular remodelling of brown adipocytes during cold-acclimation, consistent with the conclusion that metabolites could primarily drive the acute-phase response in BAT. Reporter metabolites are thought to represent key regulatory areas within a metabolic network that maintain energy homeostasis in response to environmental stressors [67]. Alterations in the abundance of N-acetylated amino acids have been identified in cold-exposed BAT [18,19,68]. Our finding that N-acetylated amino acids rapidly decrease in abundance during cold deacclimation further supports their role as potential reporter metabolite candidates.

While fatty acids are a major source of acetyl-CoA during cold exposure, M+2 labelling observed in [13-C]-glucose tracing experiments in cold-exposed mice suggests that the acetyl group of the N-acetyl-amino acids is mainly derived from glucose via glycolysis and pyruvate dehydrogenase activity [19]. N-acyl amino acids, which have a fatty acid acyl moiety rather than an acetyl group have been shown to activate UCP1 and uncouple mitochondrial respiration in a similar manner as conventional fatty acids [69,70]. In contrast, while N-acetylated amino acids have been identified in cold-exposed BAT [18,19,68], their role in thermogenesis is relatively unexplored.

High levels of N-acetylated amino acids during cold exposure are hypothesized to result from the proteolytic degradation of N-terminal acetylated proteins [19]. Our findings that protein lysine acetylation and protein content decrease in BAT during cold deacclimation are consistent with the conclusion that cold deacclimation increases proteolytic degradation of proteins, giving rise to the observed increase in N-acetyllysine abundance. Instead, our data demonstrating an increase in aminoacylase 1 expression and decreased Nat8l expression suggest that N-acetyltransferase activity is high during cold exposure, and decreases upon returning to thermoneutrality. Skeletal muscle uses a similar L-carnitine/Carnitine acetyltransferase (CrAT) buffering system for mitochondrial acetyl-CoA, particularly during times of high nutrient availability [71]. Thus, our data are consistent with the idea that N-acetyl amino acids may act as transport molecules to buffer high acetyl-CoA levels that can be converted to acetate for lipid synthesis [35] during cold acclimation. Moreover, as demonstrated for N-acetyl aspartate in brown adipocytes, the catabolism N-acetyl amino acids may provide cytosolic acetyl-CoA, which may regulate histone acetylation during thermogenesis [72]. Alternatively, N-acetyl amino acids may act as signaling molecules for organ cross-talk, as high levels of N-acetylglutamate and N-acetylaspartate have been observed in extracellular fluid derived from BAT [68].

Individual N-acetylated amino acids may play specific roles in BAT during thermogenesis. For example, N-acetyl aspartate promotes lipid turnover, and can induce UCP1 expression in a PPARα-dependent manner [73]. N-acetyl glutamate is a precursor for arginine biosynthesis and acts as an allosteric activator of the rate-limiting urea cycle enzyme, carbamoyl phosphate synthetase 1 (CPS1) [74]. Mechanistic studies have demonstrated that N-acetyl glutamate and N-acetyl methionine can impair mitochondrial membrane potential (ΔΨm) and induce mitochondrial swelling in calcium (Ca^2+^)-loaded mitochondria in a dose-dependent manner, indicating they may play a role in inducing the mitochondrial permeability transition pore (MPTP) opening [75]. N-acetyl glutamate can inhibit glutamate oxidation by glutamate dehydrogenase (GDH) as well as isocitrate dehydrogenase activity (IDH)[76]. However, the specific roles of N-acetyl amino acids in BAT during thermogenesis warrants further exploration.

In conclusion, our findings outline the progressive changes in BAT structure and function occurring in the transition from a cold acclimated state to a thermoneutral environment. Specifically, high metabolic rates and food intake during the cold acclimated state rapidly decrease in a thermoneutral environment. More gradual were the progressive declines in BAT mitochondrial uncoupling capacity, protein content, and UCP1 expression, as well as decreases in mitochondrial content, confirmed by mtDNA/nDNA ratios and electron microscopy. Metabolomic profiling showed decreased N-acetylated amino acids and major alterations in pathways involving amino acid metabolism, purine nucleotides, and the TCA cycle. Enhanced gene expression of ACY1 and ACY3 suggest that enhanced aminoacylase activity may contribute to the marked decrease in N-acetylated metabolites during cold deacclimation. Together, these findings highlight the biochemical and metabolic processes involved in BAT thermogenesis and deactivation.

## Methods

### Animals

All experimental procedures were approved by the University of Ottawa Animal Care Committee and conducted in accordance with the guidelines and principles of the Canadian Council of Animal Care. Mice were maintained on a C57BL/6J background (Jackson Laboratories) and housed in ventilated cages in a temperature-controlled room (22°C) on a 12/12 h light-dark cycle, with *ad libitum* access to a standard chow diet (44.2% carbohydrate, 6.2% fat, and 18.6% crude protein) and water. Prior to cold exposure, mice were individually housed for at least 3 days. Mice (7-9 weeks old) were acclimated to the cold (4°C) for 7 days and subsequently deacclimated at 09:00 to thermoneutrality (30°C) where they remained for 0 hours (cold-acclimated controls), or for 3, 12, 24, or 48 hours of deacclimation. Unless otherwise stated, mice were sacrificed by cervical dislocation, and interscapular BAT (iBAT) was collected for subsequent analyses.

### Metabolic phenotyping

A 12-chamber comprehensive lab animal monitoring system (CLAMS; Columbus Instruments, Columbus, OH) was used to measure volitional activity, food intake, and whole-body energy metabolism. Mice were acclimated to 4°C for 3 days prior to being transferred to individual CLAMS chambers. To minimize stress associated with a different cage environment, mice were further acclimated for 4 more days to the CLAMS chambers maintained at 4°C with *ad libitum* access to their food prior to data collection during the standard light-dark cycle. Data were collected at 4°C for 2 days prior to increasing the temperature to 30°C for 4 days, with measurements recorded every 26 minutes. For activity measurements, ambulatory activity was calculated as the sum of ambulatory beam breaks in the x, y, and Z planes (XAMB, YAMB, ZAMB). Total activity was calculated as the sum of all beam breaks detected in the X, Y, and Z dimensions (XTOT, YTOT, ZTOT).

### High resolution respirometry

Oxygen consumption was quantified in BAT using high-resolution respirometry (O2K, Oroboros Instruments, Innsbruck, Austria). Freshly isolated iBAT was minced and subsequently placed in ice-cold BIOPS buffer [BIOPS (in mM): 2.77 CaK_2_EGTA, 7.23 K_2_EGTA, 5.77 Na_2_ATP, 50 MES, 20 imidazole, 20 taurine, 15 PCr, 0.5 DTT, 6.56 MgCL_2_–6H_2_O, pH 7.1 at 0°C] and permeabilized with 50μg/ml saponin for 30 min at 4°C under gentle agitation. Samples were then washed for 30 min in ice-cold mitochondrial respiration medium [MiR05 (in mM): 110 sucrose, 60 K-lactobionate, 20 HEPES, 20 taurine, 10 KH2PO4, 3 MgCl2, 0.5 EGTA, and 1 mg/ml BSA, pH 7.1 at 37°C]. Experiments were performed in duplicate at 37 °C in 2 mL of mitochondrial respiration media. To evaluate uncoupled respiration, assays were performed in the presence of 5 µM oligomycin and 0.1 mM octanoylcarnitine to inhibit ATP synthase and activate uncoupled respiration. The assay protocol consisted of consecutive additions of 2 mM malate, 5 mM pyruvate, 10 mM glutamate (CI Leak), 10 mM succinate (CI+II Leak), 5 mM glycerol 3-phosphate (CI+II+cGpDH Leak) and Antimycin A (non-mitochondrial respiration).

### Transmission electron microscopy (TEM) of BAT

Samples for TEM were processed and imaged by the Facility for Electron Microscopy Research at McGill University. Briefly, freshly dissected iBAT was fixed with 2.5% glutaraldehyde in 0.1 M sodium cacodylate buffer (pH 7.4). The tissue was washed for 3 × 10 minutes in 0.0.1 M sodium cacodylate buffer and post-fixed in 1% aqueous OsO4 +1.5% aqueous potassium ferrocyanide for 2 hours at 4°C. The samples were dehydrated using increasing ethanol concentrations, infiltrated with graded EPON:ethanol mixtures, and embedded in 100% Epon resin. Sections were prepared using Leica Microsystems UC6 Ultramicrotome and counterstained with uranyl acetate and Reynold’s lead. TEM grids were imaged by an FEI Tecnai G2 Spirit 120 kV cryo-TEM equipped with an AMT NanoSprint15 MK2 CMOS Camera.

Images from 5 micrographs/mouse were blinded for analysis. Mitochondria were semi-automatedly segmented and annotated using the Empanada-Napari plugin [77], by training the MitoNet deep learning model. Training patches were selected at random from the images using the built-in model MitoNet_v1 to run 2D inference, and the resulting segmentations were then manually corrected in Napari. MitoNet_v1 was trained on the proofread patches for 100 iterations to generate a fine-tuned model. Outcomes from the fine-tuned model were manually checked to correct errors to ensure accurate annotation. The number and surface area of mitochondria were measured using the napari-skimage-regionprops plugin. Analysis of lipid droplet size and number were blinded and manually performed in ImageJ.

### Protein extraction and immunoblotting

Frozen iBAT was homogenized in ice-cold RIPA buffer (Millipore) supplemented with a protease inhibitor cocktail (Sigma) and Phosphatase inhibitor cocktail (Thermofisher) using a bead mill homogenizer (Fisherbrand). The homogenates were centrifuged at 10,000 g for 10 min at 4 °C, and the resulting top solidified lipid layer was discarded and the infranatant layer was carefully collected and centrifuged again at 10,000 g for 10 min at 4 °C to remove any remaining lipids. Protein concentration was determined using a commercially available bicinchoninic acid kit per the manufacturer’s protocol (ThermoFisher). Proteins were separated by SDS-PAGE under reducing conditions and transferred to PVDF membranes and incubated overnight with primary antibodies against UCP1 (1:6000; Sigma Aldrich, #U6382) and acetylated lysine-containing proteins (1:2000, Sigma Aldrich, #ST1027). Vinculin (1:5000, Abcam, ab129002) was used as a loading control. Protein bands were visualized using the ChemiDoc™ MP Imaging System (Bio-Rad) and densitometry band analyses were performed using ImageJ (NIH).

### Citrate synthase activity

Citrate synthase activity was determined in iBAT protein lysates as previously described [78]. The change in the rate of absorbance at 412 nm and pathlength was measured using a BioTek Synergy Mx Microplate Reader (BioTek Instruments) in the presence of 50 mM Tris-HCl (pH 8.0), with 0.2 mM DTNB, 0.1 mM acetyl-CoA and 0.25 mM oxaloacetate. Enzyme activity was calculated using the extinction coefficient of 13.6 mM^−1^cm^−1^ for citrate synthase.

### Mitochondrial and nuclear DNA quantification

DNA was extracted from frozen iBAT as previously described [79]. SsoAdvanced™ Universal SYBR® Green Supermix (Bio-Rad) was used for quantitative PCR on the CFX96 Real-Time PCR Detection System (Bio-Rad). Mitochondrial DNA (mtDNA) to nuclear DNA (nDNA) ratios were determined through qPCR against mitochondrial DNA gene mt-ND1 (FWD: 5′-CTAGCAGAAACAAACCGGGC-3′, REV: 5′-CCGGCTGCGTATTCTACGTT-3′) and nuclear DNA gene HK2 (FWD: 5′-GCCAGCCTCTCCTGATTTTAGTGT-3′, REV: 5′-GGGAACACAAAAGACCTCTTCTGG-3′).

### RNA extraction and quantitative PCR

RNA was extracted from frozen iBAT using Trizol (ThermoFisher, #15596026) according to the manufacturer’s instructions. Total RNA concentration was measured using a NanoDrop™ 2000 UV–Vis spectrophotometer (Thermo Scientific). cDNA was synthesized using the All-In-One 5X RT MasterMix (ABM, #G592). Quantitative PCR was performed using the SsoAdvanced Universal SYBR Green Supermix (Bio-Rad, #1725272) and run on the CFX96 (Bio-Rad). Primer pairs for target genes are outlined in **Table 1**. As a control for between-sample variability, mRNA levels were normalized to the geometric mean of beta-2 microglobulin (B2M), hypoxanthine phosphoribosyltransferase 1 (Hrpt1), glyceraldehyde-3-phosphate dehydrogenase (GAPDH). Relative transcript expression was calculated using the 2^−ΔΔCt^ method [80].

**Table 1:**
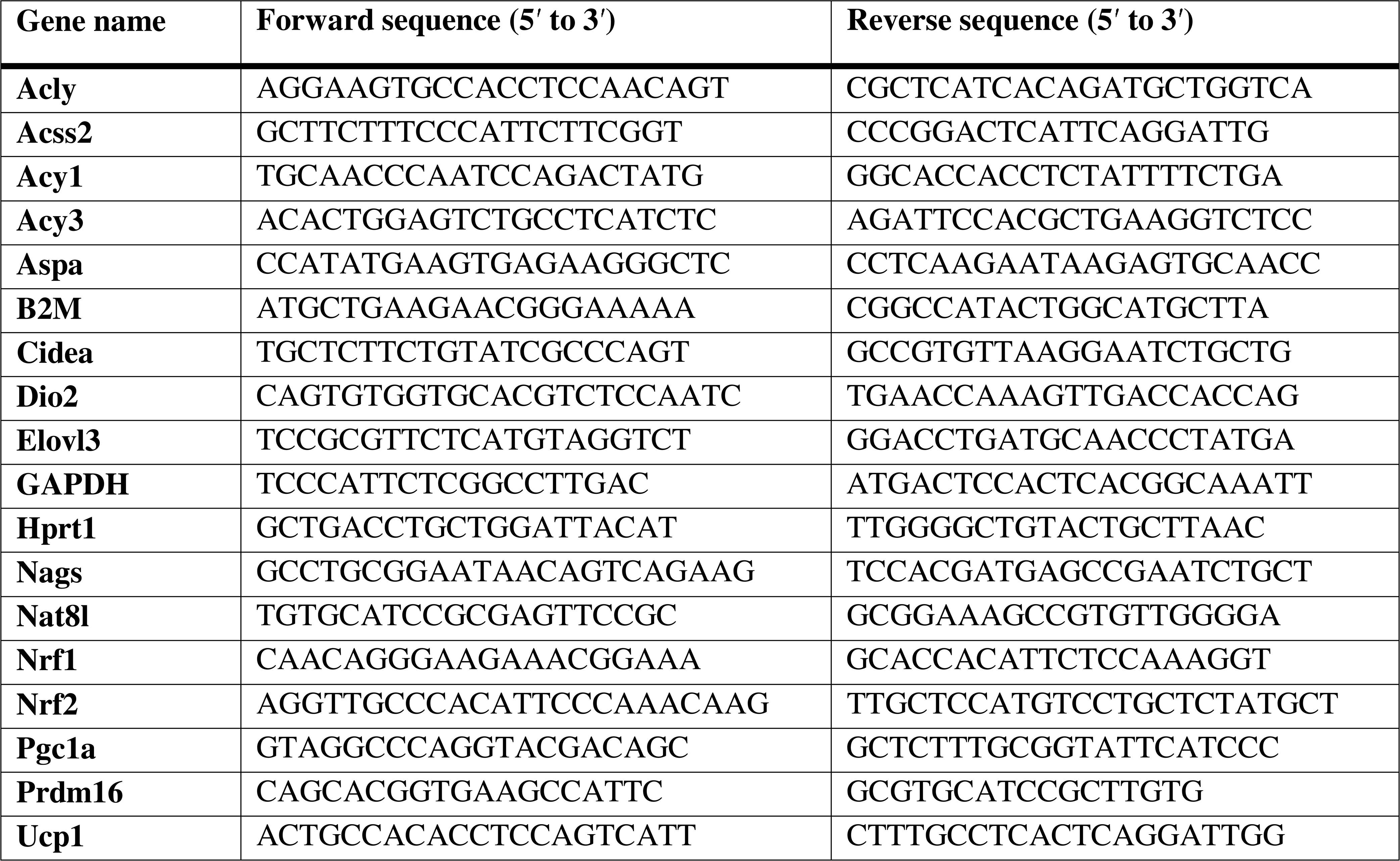
Primer sequences used for quantitative PCR.

### Targeted metabolomics

Frozen iBAT was homogenized using a bead mill homogenizer at 4 °C (Fisherbrand Bead Mill 24 Homogenizer) in a −20 °C equilibrated solution containing methanol, water, and acetonitrile (OmniSolv, Sigma). Homogenates were then incubated with a 2:1 dichloromethane:water solution on ice for 10 min. The polar and non-polar phases were separated by centrifugation at 3200 x g for 10 min at 1 °C. The upper polar phase was evapourated using a refrigerated CentriVap Vacuum Concentrator at −4 °C (LabConco Corporation, Kansas City, MO). Samples were run on an Agilent 6470A tandem quadrupole mass spectrometer equipped with a 1290 Infinity II ultra-high performance LC (Agilent Technologies) utilizing the Metabolomics Dynamic MRM Database and Method (Agilent), using ion-pairing reverse phase chromatography[81]. This method was further optimized for phosphate-containing metabolites with the addition of 5 μM InfinityLab deactivator (Agilent Technologies) to mobile phases A and B, which requires decreasing the backflush acetonitrile to 90%. Multiple reaction monitoring (MRM) transitions were optimized using authentic standards and quality control samples. Metabolites were quantified by integrating the area under the curve of each compound using external standard calibration curves with Mass Hunter Quant (Agilent). No corrections for ion suppression or enhancement were performed; as such, uncorrected metabolite concentrations are presented. Further data processing analysis was conducted in MetaboAnalyst 6.0 [82]. Metabolite set enrichment analysis (MSEA) was performed using over-representation analysis with the Small Molecule Pathway Database as a library.

### Data mining and network analysis

Data mining and network analyses were performed using in-house software written in Matlab 2024b (Matworks Inc) and https://complimet.ca/sidco (59). ReliefF was used for feature selection (command relief running under Matlab) using Euclidean distances as a metric for feature similarity and the 200 nearest neighbors for weight assessment. Correlation analysis using the distance correlation calculations were performed using SIDCO (58) and in-house routines developed in Matlab. Correlation p-values were calculated using Student’s t cumulative distribution function. Metabolite differences between groups were determined using linear regression comparisons of correlation values in the two groups for each metabolite, as previously described (59).

### Bioinformatic analyses of iBAT transcriptomics and proteomics

Publicly available published datasets from BAT cold exposure and cold acclimation studies were identified for analyses of aminoacylase and N-acetyltransferase expression. Analysed transcriptomic data from mice housed at thermoneutrality or cold (4°C) for 4 weeks [37] were obtained from Sanchez et al (2018) supplementary files. Proteomics data were obtained from Cutler *et al*. (2025), where mice were housed under thermoneutral (30°C) or cold (5°C) conditions for 3 weeks [38]. To identify proteins involved in the (de)acetylation of proteins and amino acids during cold acclimation, the GO terms (https://www.ebi.ac.uk/QuickGO/) N-acetyltransferase activity (GO:0008080) and aminoacylase activity (GO:0004046) were used to select targets with an FDR<0.05 as computed by the authors.

### Statistical analysis

Statistical analyses were performed using GraphPad Prism 10 (GraphPad Prism, La Jolla, CA, USA) and all values are reported as mean ± SD with a significance level of *p* < 0.05. For metabolic phenotyping analyses performed with CLAMS, a one-way repeated measures (RM) ANOVA was used to determine the effects cold-deacclimation. For subsequent experiments using cold acclimated (4°C) and cold deacclimated (30°C) mice at varying timepoints (3, 12, 24, or 48 hours of deacclimation), a one-way ANOVA was used to determine the effects cold-deacclimation. Tukey posthoc-tests and multiple comparisons tests were used to assess changes between the timepoints of cold-deacclimated state. For metabolomic comparisons between cold acclimated and 48 h cold deacclimated mice, a two-tailed Student’s t-test was used.

## Abbreviation list

ACC1: acetyl-CoA carboxylase
ACLY: ATP-citrate lyase
ACY1: aminoacylase 1
ACY3: aminoacylase3
ACSS2: acetyl-CoA synthetase 2
ASPA: aspartoacylase
BAT: brown adipose tissue
CIDEA: cell death inducing DFFA like effector a
CLAMS: comprehensive lab animal monitoring system
CRAT: carnitine acetyltransferase
DIO2: type 2 iodothyronine deiodinase
ELOVL3: ELOVL fatty acid elongase 3
FASN: fatty acid synthase
FFA: free fatty acid
G3P: glycerol-3-phosphate
GDH: glutamate dehydrogenase
GSH: glutathione
GSSG: oxidized glutathione
HRR: high-resolution respirometry
IDH: isocitrate dehydrogenase
LCMS: liquid chromatography-mass spectroscopy
mGDPDH: mitochondrial glycerol-3-phosphate dehydrogenase
MPTP: mitochondrial permeability transition pore
MSEA: metabolite set enrichment analysis
NAGS: N-acetylglutamate synthase
NAT8L: N-acetyltransferase 8-like
NRF1: nuclear respiratory factor 1
NRF2: nuclear respiratory factor 2
NST: non-shivering thermogenesis
PANK1: pantothenate kinase 1
PGC1α: peroxisome proliferator-activated receptor γ coactivator-1α
PKA: protein kinase A
PRDM16: PR domain containing 16
RER: respiratory exchange ratio
ROS: reactive oxygen species
SNS: sympathetic nervous system
TCA cycle: tricarboxylic acid cycle
TEM: transmission electron microscopy
UCP1: uncoupling protein 1

## Acknowledgements

The authors would like to thank Jian Ying Xuan for her excellent technical support, the Metabolomics Core, Cell Biology and Image Acquisition Core, Animal Care and Veterinary Services Behavioural Core, and the Histology Core, all funded by the University of Ottawa, Ottawa, Canada. The authors would also like to thank Kelly Sears, Hojatollah Vali, and Jeannie Mui at the Facility for Electron Microscopy Research of McGill University for help in microscope operation and data collection. This research was funded through a Discovery grant from the Natural Sciences and Engineering Research Council (NSERC) of Canada.

## Conflict of Interest

The authors declare no conflicts of interest.

## Availability of Data and Materials

All data generated for this manuscript are presented in the main manuscript or additional supporting files.

## Author Contributions

Conceived and designed the experiments: CAP, MEH

Performed experiments: CAP, NK, EM, LH, LSK, VV, MK, ZEH, YB

Analyzed the data: CAP, NK, EM, LH, LSK, VV, MK, ZEH, MCC

Wrote the paper: CAP

Reviewed and edited: CAP, NK, EM, LH, LSK, VV, MK, ZEH, YB, MCC, MEH

## Supplementary Legends

**Table S1:** Published RNA-seq [37] and proteomics [38] datasets identifying proteins involved in the (de)acetylation of proteins and amino acids during cold acclimation using the GO terms N-acetyltransferase activity (GO:0008080) and aminoacylase activity (GO:0004046).

**Figure S1:**
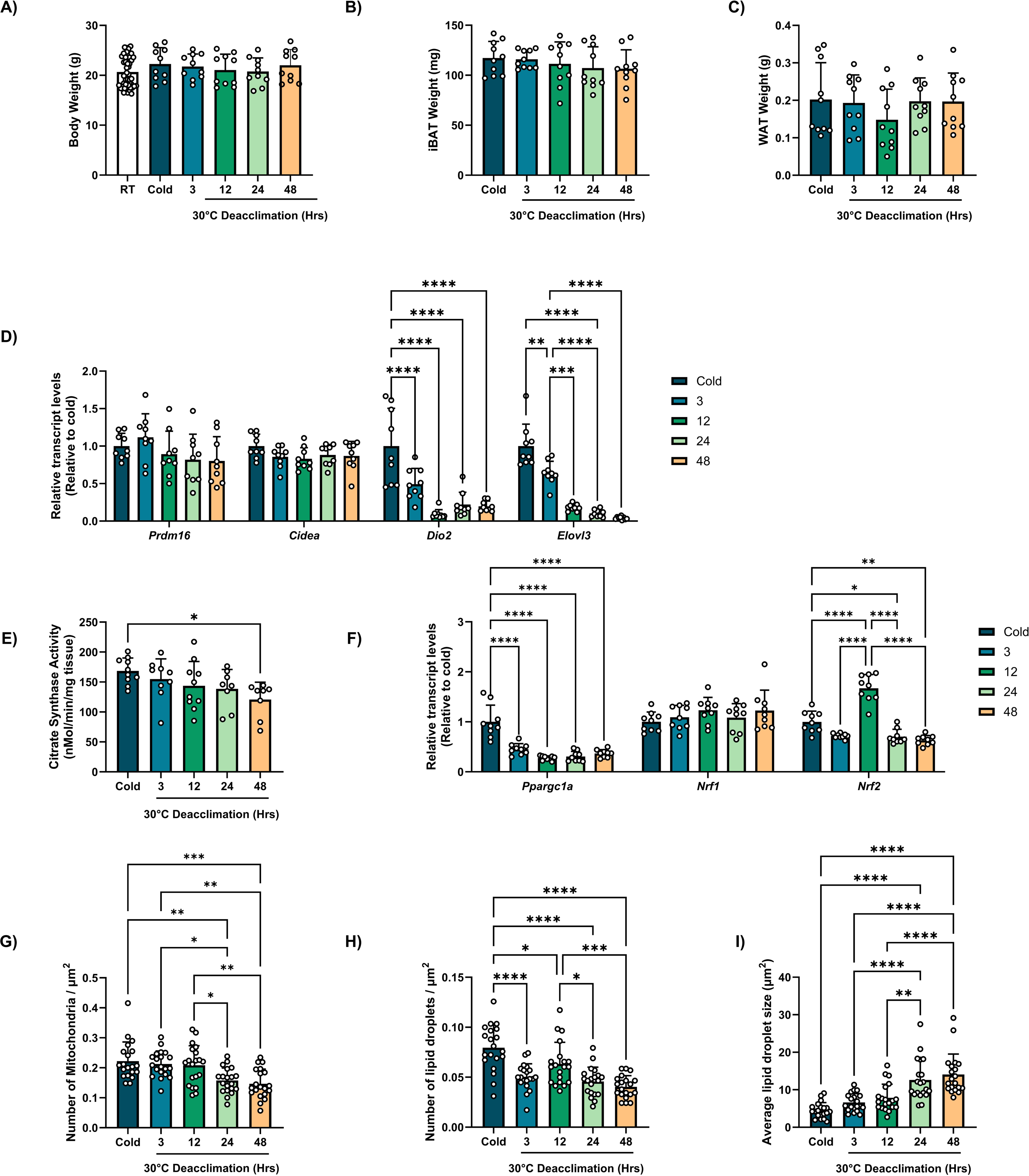
**(A)** Body weights of mice prior to the cold acclimation period (RT). Mice were then acclimated to the cold (4°C) for 7 days, and subsequently transferred to thermoneutrality (30°C) for 3 h, 12 h, 24 h, or 48 h, (n=10/group). **(B)** Brown adipose tissue weight, (n=10/group). **(C)** White adipose tissue (epididymal) weight, (n=10/group). **(D)** qPCR was used to determine the relative transcript levels of BAT activation markers, PR domain containing 16 (*Prdm16*), cell death inducing DFFA like effector a (*Cidea*), type 2 iodothyronine deiodinase (*Dio2*), and ELOVL fatty acid elongase 3 (*Elvol3*), (n=9/group). **(E)** Citrate synthase activity, (n=8-10/group). **(F)** qPCR was used to determine the relative transcript levels of mitochondrial biogenesis transcription factors including PGC1α (Ppargc1a), nuclear factor 1 (NRF) and 2 (NRF2), (n=9/group). **(G-I)** TEM images were analyzed by quantitative morphometry for **(G)** Number of mitochondria, **(H)** number of lipid droplets, and **(I)** average lipid droplet size, (n=5 TEM images from 4 mice/group). See also Figure 2. Comparisons between timepoints were determined using a one-way ANOVA with Tukey post-hoc tests, *<p<0.05 **p<0.01, ***p<0.001, ****p<0.0001. All values are presented as means ± SD.

**Figure S2.**
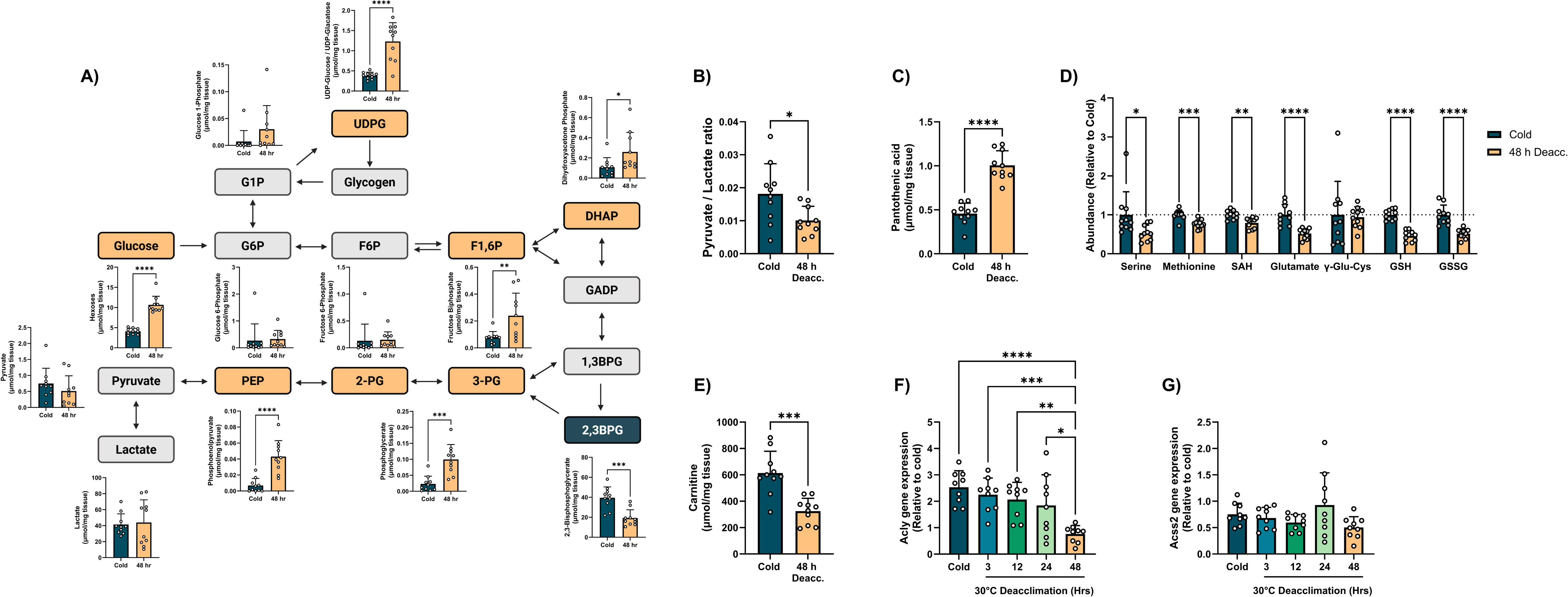
**(A)** Quantitative analysis of individual metabolites relating to glycolysis, (n=10/group). **(B)** The ratio of pyruvate to lactate, (n=10/group). **(C)** Quantitative abundance of pantothenate, (n=10/group). **(D)** The abundances of serine, methionine, S-adenosyl homocysteine (SAH), glutamate, gamma-glu-cys, oxidized glutathione (GSSG) and reduced glutathione (GSH) (relative to cold), (n=10/group). **(E)** Quantitative abundance of carnitine, (n=10/group). **(G)(F-G)** qPCR of gene expression of **(F)** ATP citrate lyase and **(G)** acetyl-CoA synthetase 2, (n=9/group). Comparisons between timepoints were determined using a one-way ANOVA with Tukey post-hoc tests. Comparisons for individual metabolites were determined using a two-tailed Student’s t-test. All values are presented as means ± SD. *<p<0.05 **p<0.01, ***p<0.001, ****p<0.0001.

## Notes

**Grant support:** Natural Sciences and Engineering Research Council of Canada (MH)

### Competing Interest Statement

The authors have declared no competing interest.

## References

[1] D.P. Blondin, S.M. Labbé, H.C. Tingelstad, C. Noll, M. Kunach, S. Phoenix, B. Guérin, É.E. Turcotte, A.C. Carpentier, D. Richard, F. Haman, Increased Brown Adipose Tissue Oxidative Capacity in Cold-Acclimated Humans, J Clin Endocrinol Metab 99 (2014) E438–E446. 10.1210/JC.2013-3901.

[2] D.G. Nicholls, R.M. Locke, Thermogenic mechanisms in brown fat., Physiol Rev 64 (1984) 1–64. 10.1152/physrev.1984.64.1.1.

[3] J. Himms-Hagen, Brown adipose tissue metabolism and thermogenesis., Annu Rev Nutr 5 (1985) 69–94. 10.1146/annurev.nu.05.070185.000441.

[4] G.M. Heaton, R.J. Wagenvoord, A. Kemp, D.G. Nicholls, Brown-Adipose-Tissue Mitochondria: Photoaffinity Labelling of the Regulatory Site of Energy Dissipation, Eur J Biochem 82 (1978) 515–521. 10.1111/J.1432-1033.1978.TB12045.X.

[5] S. Enerbäck, A. Jacobsson, E.M. Simpson, C. Guerra, H. Yamashita, M.E. Harper, L.P. Kozak, Mice lacking mitochondrial uncoupling protein are cold-sensitive but not obese, Nature 1997 387:6628 387 (1997) 90–94. 10.1038/387090a0.

[6] B. Cannon, J.A.N. Nedergaard, Brown adipose tissue: function and physiological significance, Physiol Rev 84 (2004) 277–359. 10.1152/physrev.00015.2003.

[7] V. Golozoubova, B. Cannon, J. Nedergaard, UCP1 is essential for adaptive adrenergic nonshivering thermogenesis, Am J Physiol Endocrinol Metab 291 (2006). 10.1152/ajpendo.00387.2005.

[8] J. Himms Hagen, The role of brown adipose tissue in the calorigenic effect of adrenaline and noradrenaline in cold-acclimated rats, J Physiol 205 (1969) 393. 10.1113/jphysiol.1969.sp008973.

[9] S. Collins, R.S. Surwit, The beta-adrenergic receptors and the control of adipose tissue metabolism and thermogenesis., Recent Prog Horm Res 56 (2001) 309–328. 10.1210/rp.56.1.309.

[10] M. Uldry, W. Yang, J. St-Pierre, J. Lin, P. Seale, B.M. Spiegelman, Complementary action of the PGC-1 coactivators in mitochondrial biogenesis and brown fat differentiation, Cell Metab 3 (2006) 333–341. 10.1016/j.cmet.2006.04.002.

[11] P. Puigserver, Z. Wu, C.W. Park, R. Graves, M. Wright, B.M. Spiegelman, A cold-inducible coactivator of nuclear receptors linked to adaptive thermogenesis, Cell 92 (1998) 829–839. 10.1016/s0092-8674(00)81410-5.

[12] L.A. de Jesus, S.D. Carvalho, M.O. Ribeiro, M. Schneider, S.-W. Kim, J.W. Harney, P.R. Larsen, A.C. Bianco, The type 2 iodothyronine deiodinase is essential for adaptive thermogenesis in brown adipose tissue, Journal of Clinical Investigation 108 (2001) 1379. 10.1172/JCI13803.

[13] J.E. Silva, P.R. Larsen, Adrenergic activation of triiodothyronine production in brown adipose tissue, Nature 305 (1983) 712–713. 10.1038/305712a0.

[14] A. Guilherme, B. Yenilmez, A.H. Bedard, F. Henriques, D. Liu, A. Lee, L. Goldstein, M. Kelly, S.M. Nicoloro, M. Chen, L. Weinstein, S. Collins, M.P. Czech, Control of Adipocyte Thermogenesis and Lipogenesis through β3-Adrenergic and Thyroid Hormone Signal Integration, Cell Rep 31 (2020) 107598. 10.1016/j.celrep.2020.107598.

[15] L. Bukowiecki, A.J. Collet, N. Follea, Brown adipose tissue hyperplasia: a fundamental mechanism of adaptation to cold and hyperphagia, Am J Physiol 242 (1982). 10.1152/ajpendo.1982.2426.E353.

[16] I.L. Cameron, R.E. Smith, Cytological Responses of Brown Fat Tissue in Cold-Exposed Rats, Journal of Cell Biology 23 (1964) 89–100. 10.1083/jcb.23.1.89.

[17] R.A. Piñol, A.S. Mogul, C.K. Hadley, A. Saha, C. Li, V. Škop, H.S. Province, C. Xiao, O. Gavrilova, M.J. Krashes, M.L. Reitman, Preoptic BRS3 neurons increase body temperature and heart rate via multiple pathways, Cell Metab 33 (2021) 1389–1403.e6. 10.1016/j.cmet.2021.05.001.

[18] E.L. Mills, K.A. Pierce, M.P. Jedrychowski, R. Garrity, S. Winther, S. Vidoni, T. Yoneshiro, J.B. Spinelli, G.Z. Lu, L. Kazak, A.S. Banks, M.C. Haigis, S. Kajimura, M.P. Murphy, S.P. Gygi, C.B. Clish, E.T. Chouchani, Accumulation of succinate controls activation of adipose tissue thermogenesis, Nature 560 (2018) 102–106. 10.1038/s41586-018-0353-2.

[19] S.M. Jung, W.G. Doxsey, J. Le, J.A. Haley, L. Mazuecos, A.K. Luciano, H. Li, C. Jang, D.A. Guertin, In vivo isotope tracing reveals the versatility of glucose as a brown adipose tissue substrate, Cell Rep 36 (2021) 109459. 10.1016/j.celrep.2021.109459.

[20] C.J. Gordon, Relationship between autonomic and behavioral thermoregulation in the mouse, Physiol Behav 34 (1985) 687–690. 10.1016/0031-9384(85)90365-8.

[21] B. Cannon, J. Nedergaard, Nonshivering thermogenesis and its adequate measurement in metabolic studies, J Exp Biol 214 (2011) 242–253. 10.1242/jeb.050989.

[22] J.R. Brobeck, Food Intake as a Mechanism of Temperature Regulation, Yale J Biol Med 20 (1948) 545. https://pmc.ncbi.nlm.nih.gov/articles/PMC2602369/ (accessed October 30, 2024).

[23] N.C. Bal, S. Singh, F.C.G. Reis, S.K. Maurya, S. Pani, L.A. Rowland, M. Periasamy, Both brown adipose tissue and skeletal muscle thermogenesis processes are activated during mild to severe cold adaptation in mice, Journal of Biological Chemistry 292 (2017) 16616–16625. 10.1074/jbc.M117.790451.

[24] K. Dümmler, S. Müller, H.J. Seitz, Regulation of adenine nucleotide translocase and glycerol 3-phosphate dehydrogenase expression by thyroid hormones in different rat tissues, Biochemical Journal 317 (1996) 913–918. 10.1042/BJ3170913.

[25] K.I. Ohkawa, M.T. Vogt, E. Farber, Unusually high mitochondrial alpha glycerophosphate dehydrogenase activity in rat brown adipose tissue, J Cell Biol 41 (1969) 441. 10.1083/jcb.41.2.441.

[26] A.A. Wendel, T.M. Lewin, R.A. Coleman, Glycerol-3-phosphate acyltransferases: Rate limiting enzymes of triacylglycerol biosynthesis, Biochimica et Biophysica Acta (BBA) - Molecular and Cell Biology of Lipids 1791 (2009) 501–506. 10.1016/j.bbalip.2008.10.010.

[27] P. Puigserver, D. Herron, M. Gianotti, A. Palou, B. Cannon, J. Nedergaard, Induction and degradation of the uncoupling protein thermogenin in brown adipocytes in vitro and in vivo. Evidence for a rapidly degradable pool, Biochem J 284 (Pt 2) (1992) 393–398. 10.1042/BJ2840393.

[28] A. Bartelt, O.T. Bruns, R. Reimer, H. Hohenberg, H. Ittrich, K. Peldschus, M.G. Kaul, U.I. Tromsdorf, H. Weller, C. Waurisch, A. Eychmüller, P.L.S.M. Gordts, F. Rinninger, K. Bruegelmann, B. Freund, P. Nielsen, M. Merkel, J. Heeren, Brown adipose tissue activity controls triglyceride clearance, Nat Med 17 (2011) 200–205. 10.1038/nm.2297.

[29] H. Zhou, H. Zhang, R. Ye, C. Yan, J. Lin, Y. Huang, X. Jiang, S. Yuan, L. Chen, R. Jiang, K. Zheng, Z. Cheng, Z. Zhang, M. Dong, W. Jin, Pantothenate protects against obesity via brown adipose tissue activation, Am J Physiol Endocrinol Metab 323 (2022) E69–E79. 10.1152/ajpendo.00293.2021.

[30] Y. Takeda, P. Dai, Functional roles of pantothenic acid, riboflavin, thiamine, and choline in adipocyte browning in chemically induced human brown adipocytes, Sci Rep 14 (2024) 1–15. 10.1038/s41598-024-69364-w.

[31] M. U Din, T. Saari, J. Raiko, N. Kudomi, S.F. Maurer, M. Lahesmaa, T. Fromme, E.Z. Amri, M. Klingenspor, O. Solin, P. Nuutila, K.A. Virtanen, Postprandial Oxidative Metabolism of Human Brown Fat Indicates Thermogenesis, Cell Metab 28 (2018) 207–216.e3. 10.1016/j.cmet.2018.05.020.

[32] A. Perdikari, G.G. Leparc, M. Balaz, N.D. Pires, M.E. Lidell, W. Sun, F. Fernandez-Albert, S. Müller, N. Akchiche, H. Dong, L. Balazova, L. Opitz, E. Röder, H. Klein, P. Stefanicka, L. Varga, P. Nuutila, K.A. Virtanen, T. Niemi, M. Taittonen, G. Rudofsky, J. Ukropec, S. Enerbäck, E. Stupka, H. Neubauer, C. Wolfrum, BATLAS: Deconvoluting Brown Adipose Tissue, Cell Rep 25 (2018) 784–797.e4. 10.1016/j.celrep.2018.09.044.

[33] R.J. Mailloux, C.N.-K. Adjeitey, J.Y. Xuan, M.-E. Harper, Crucial yet divergent roles of mitochondrial redox state in skeletal muscle vs. brown adipose tissue energetics, The FASEB Journal 26 (2012) 363–375. 10.1096/fj.11-189639.

[34] S. Zhao, A.M. Torres, R.A. Henry, S. Trefely, M. Wallace, J. V. Lee, A. Carrer, A. Sengupta, S.L. Campbell, Y.M. Kuo, A.J. Frey, N. Meurs, J.M. Viola, I.A. Blair, A.M. Weljie, C.M. Metallo, N.W. Snyder, A.J. Andrews, K.E. Wellen, ATP-Citrate Lyase Controls a Glucose-to-Acetate Metabolic Switch, Cell Rep 17 (2016) 1037–1052. 10.1016/j.celrep.2016.09.069.

[35] E.D. Korobkina, C.M. Calejman, J.A. Haley, M.E. Kelly, H. Li, M. Gaughan, Q. Chen, H.L. Pepper, H. Ahmad, A. Boucher, S.M. Fluharty, T.Y. Lin, A. Lotun, J. Peura, S. Trefely, C.R. Green, P. Vo, C.F. Semenkovich, J.R. Pitarresi, J.B. Spinelli, O. Aydemir, C.M. Metallo, M.D. Lynes, C. Jang, N.W. Snyder, K.E. Wellen, D.A. Guertin, Brown fat ATP-citrate lyase links carbohydrate availability to thermogenesis and guards against metabolic stress, Nat Metab (2024). 10.1038/S42255-024-01143-3.

[36] M.W. Anders, W. Dekant, Aminoacylases, Adv Pharmacol 27 (1994) 431–448. 10.1016/s1054-3589(08)61042-x.

[37] J. Sanchez-Gurmaches, Y. Tang, N.Z. Jespersen, M. Wallace, C. Martinez Calejman, S. Gujja, H. Li, Y.J.K. Edwards, C. Wolfrum, C.M. Metallo, S. Nielsen, C. Scheele, D.A. Guertin, Brown Fat AKT2 Is a Cold-Induced Kinase that Stimulates ChREBP-Mediated De Novo Lipogenesis to Optimize Fuel Storage and Thermogenesis, Cell Metab 27 (2018) 195–209.e6. 10.1016/j.cmet.2017.10.008.

[38] H.B. Cutler, S. Jall-Rogg, S. Thillainadesan, K.C. Cooke, S.W.C. Masson, J.M. Sligar, J.G. Crowston, L. Carroll, J. Stöckli, D.E. James, S. Madsen, Cold exposure stimulates cross-tissue metabolic rewiring to fuel glucose-dependent thermogenesis in brown adipose tissue, Science Advances 11 (2025) 7369. 10.1126/sciadv.adt7369.

[39] E.R. Suter, The fine structure of brown adipose tissue. I. Cold-induced changes in the rat, J Ultrastruct Res 26 (1969) 216–241. 10.1016/S0022-5320(69)80003-1.

[40] K. Hori, T. Ishigaki, K. Koyama, M. Kaya, J. Tsujita, S. Hori, Adaptive Changes in the Thermogenesis of Rats by Cold Acclimation and Deacclimation, Jpn J Physiol 48 (1998) 505–508. 10.2170/jjphysiol.48.505.

[41] S.M. Labbé, A. Caron, I. Bakan, M. Laplante, A.C. Carpentier, R. Lecomte, D. Richard, In vivo measurement of energy substrate contribution to cold-induced brown adipose tissue thermogenesis, The FASEB Journal 29 (2015) 2046–2058. 10.1096/fj.14-266247.

[42] S. Reichling, R.G. Ridley, H. V. Patel, C.B. Harley, K.B. Freeman, Loss of brown adipose tissue uncoupling protein mRNA on deacclimation of cold-exposed rats, Biochem Biophys Res Commun 142 (1987) 696–701. 10.1016/0006-291X(87)91470-7.

[43] G.L. McKie, H. Shamshoum, K.L. Hunt, H.H.A. Thorpe, H.A. Dibe, J.Y. Khokhar, C.A. Doucette, D.C. Wright, Intermittent cold exposure improves glucose homeostasis despite exacerbating diet-induced obesity in mice housed at thermoneutrality, J Physiol 600 (2022) 829–845. 10.1113/JP281774.

[44] Y. Ravussin, C. Xiao, O. Gavrilova, M.L. Reitman, Effect of intermittent cold exposure on brown fat activation, obesity, and energy homeostasis in mice, PLoS One 9 (2014) e85876. 10.1371/journal.pone.0085876.

[45] K.I. Stanford, R.J.W. Middelbeek, K.L. Townsend, D. An, E.B. Nygaard, K.M. Hitchcox, K.R. Markan, K. Nakano, M.F. Hirshman, Y.H. Tseng, L.J. Goodyear, Brown adipose tissue regulates glucose homeostasis and insulin sensitivity, J Clin Invest 123 (2013) 215–223. 10.1172/JCI62308.

[46] J. Wu, X. Ji, Z. Hu, B. Sun, L. Li, Physiological responses and thermal sensation in the recovery period after extremely cold exposure, Build Environ 200 (2021) 107958. 10.1016/j.buildenv.2021.107958.

[47] B. Buzadžić, A. Korać, V. Petrović, A. Vasilijević, A. Janković, B. Korać, Adaptive changes in interscapular brown adipose tissue during reacclimation after cold: The role of redox regulation, J Therm Biol 32 (2007) 261–269. 10.1016/j.jtherbio.2007.01.015.

[48] K. Hori, T. Ishigaki, K. Koyama, H. Otani, N. Kanoh, T. Tsujimura, N. Terada, S. Hori, Memory of long-term cold acclimation in deacclimated Wistar rats, J Therm Biol 31 (2006) 124–130. 10.1016/j.jtherbio.2005.11.006.

[49] B.P. Leitner, L.S. Weiner, M. Desir, P.A. Kahn, D.J. Selen, C. Tsang, G.M. Kolodny, A.M. Cypess, Kinetics of human brown adipose tissue activation and deactivation, Int J Obes 43 (2019) 633–637. 10.1038/s41366-018-0104-3.

[50] A.M.J. Claessens-Van Ooijen, K.R. Westerterp, L. Wouters, P.F.M. Schoffelen, A.A. Van Steenhoven, W.D. Van Marken Lichtenbelt, Heat Production and Body Temperature During Cooling and Rewarming in Overweight and Lean Men, Obesity 14 (2006) 1914–1920. 10.1038/OBY.2006.223.

[51] J.R. Davis, A.R. Tagliaferro, J.S. Roberts, J.O. Hill, Effects of early cold adaptation on food efficiency and dietary-induced thermogenesis in the adult rat, Physiol Behav 29 (1982) 135–140. 10.1016/0031-9384(82)90377-8.

[52] T. Scott Johnson, S. Murray, J.B. Young, L. Landsberg, Restricted food intake limits brown adipose tissue hypertrophy in cold exposure, Life Sci 30 (1982) 1423–1426. 10.1016/0024-3205(82)90555-0.

[53] J.B. Young, E. Saville, N.J. Rothwell, M.J. Stock, L. Landsberg, Effect of Diet and Cold Exposure on Norepinephrine Turnover in Brown Adipose Tissue of the Rat, J Clin Invest 69 (1982) 1061–1071. 10.1172/JCI110541.

[54] M.J. Stock, The role of brown adipose tissue in diet-induced thermogenesis, Proceedings of the Nutrition Society 48 (1989) 189–196. 10.1079/pns19890029.

[55] F.C. Hankenson, J.O. Marx, C.J. Gordon, J.M. David, Effects of Rodent Thermoregulation on Animal Models in the Research Environment, Comp Med 68 (2018) 425. 10.30802/AALAS-CM-18-000049.

[56] T. Heim, D. Hull, The blood flow and oxygen consumption of brown adipose tissue in the new-born rabbit, J Physiol 186 (1966) 42. 10.1113/jphysiol.1966.sp008019.

[57] D.O. Foster, M.L. Frydman, Tissue distribution of cold-induced thermogenesis in conscious warm- or cold-acclimated rats reevaluated from changes in tissue blood flow: the dominant role of brown adipose tissue in the replacement of shivering by nonshivering thermogenesis, Can J Physiol Pharmacol 57 (1979) 257–270. 10.1139/Y79-039.

[58] H. V. Patel, K.B. Freeman, M. Desautels, Selective loss of uncoupling protein mRNA in brown adipose tissue on deacclimation of cold-acclimated mice, Biochem Cell Biol 65 (1987) 955–959. 10.1139/O87-124.

[59] M. Desautels, J. Himms-Hagen, Parallel regression of cold-induced changes in ultrastructure, composition, and properties of brown adipose tissue mitochondria during recovery of rats from acclimation to cold, Can J Biochem 58 (1980) 1057–1068. 10.1139/O80-143.

[60] V. Farkas, G. Kelenyi, A. Sandor, A Dramatic Accumulation of Glycogen in the Brown Adipose Tissue of Rats Following Recovery from Cold Exposure, Arch Biochem Biophys 365 (1999) 54–61. 10.1006/abbi.1999.1157.

[61] P.B. Jakus, A. Sandor, T. Janaky, V. Farkas, Cooperation between BAT and WAT of rats in thermogenesis in response to cold, and the mechanism of glycogen accumulation in BAT during reacclimation, J Lipid Res 49 (2008) 332–339. 10.1194/jlr.M700316-JLR200.

[62] G. Park, J.A. Haley, J. Le, S.M. Jung, T.P. Fitzgibbons, E.D. Korobkina, H. Li, S.M. Fluharty, Q. Chen, J.B. Spinelli, C.M. Trivedi, C. Jang, D.A. Guertin, Quantitative analysis of metabolic fluxes in brown fat and skeletal muscle during thermogenesis, Nature Metabolism 2023 5:7 5 (2023) 1204–1220. 10.1038/s42255-023-00825-8.

[63] M.R. Bornstein, M.D. Neinast, X. Zeng, Q. Chu, J. Axsom, C. Thorsheim, K. Li, M.C. Blair, J.D. Rabinowitz, Z. Arany, Comprehensive quantification of metabolic flux during acute cold stress in mice, Cell Metab 35 (2023) 2077–2092.e6. 10.1016/j.cmet.2023.09.002.

[64] Z. Wang, T. Ning, A. Song, J. Rutter, Q.A. Wang, L. Jiang, Chronic cold exposure enhances glucose oxidation in brown adipose tissue, EMBO Rep 21 (2020). 10.15252/embr.202050085.

[65] X.X. Yu, D.A. Lewin, W. Forrest, S.H. Adams, Cold elicits the simultaneous induction of fatty acid synthesis and β-oxidation in murine brown adipose tissue: prediction from differential gene expression and confirmation in vivo, The FASEB Journal 16 (2002) 155–168. 10.1096/fj.01-0568com.

[66] C. Martinez Calejman, S. Trefely, S.W. Entwisle, A. Luciano, S.M. Jung, W. Hsiao, A. Torres, C.M. Hung, H. Li, N.W. Snyder, J. Villén, K.E. Wellen, D.A. Guertin, mTORC2-AKT signaling to ATP-citrate lyase drives brown adipogenesis and de novo lipogenesis, Nat Commun 11 (2020) 1–16. 10.1038/s41467-020-14430-w.

[67] K.R. Patil, J. Nielsen, Uncovering transcriptional regulation of metabolism by using metabolic network topology, Proc Natl Acad Sci U S A 102 (2005) 2685–2689. 10.1073/pnas.0406811102.

[68] A.R.P. Verkerke, D. Wang, N. Yoshida, Z.H. Taxin, X. Shi, S. Zheng, Y. Li, C. Auger, S. Oikawa, J.S. Yook, M. Granath-Panelo, W. He, G.F. Zhang, M. Matsushita, M. Saito, R.E. Gerszten, E.L. Mills, A.S. Banks, Y. Ishihama, P.J. White, R.W. McGarrah, T. Yoneshiro, S. Kajimura, BCAA-nitrogen flux in brown fat controls metabolic health independent of thermogenesis, Cell 187 (2024) 2359–2374.e18. 10.1016/j.cell.2024.03.030.

[69] J.Z. Long, K.J. Svensson, L.A. Bateman, H. Lin, T. Kamenecka, I.A. Lokurkar, J. Lou, R.R. Rao, M.R.R. Chang, M.P. Jedrychowski, J.A. Paulo, S.P. Gygi, P.R. Griffin, D.K. Nomura, B.M. Spiegelman, The Secreted Enzyme PM20D1 Regulates Lipidated Amino Acid Uncouplers of Mitochondria, Cell 166 (2016) 424–435. 10.1016/j.cell.2016.05.071.

[70] Y. Gao, I.G. Shabalina, G.R.F. Braz, B. Cannon, G. Yang, J. Nedergaard, Establishing the potency of N-acyl amino acids versus conventional fatty acids as thermogenic uncouplers in cells and mitochondria from different tissues, Biochimica et Biophysica Acta (BBA) - Bioenergetics 1863 (2022) 148542. 10.1016/j.bbabio.2022.148542.

[71] M.N. Davies, L. Kjalarsdottir, J.W. Thompson, L.G. Dubois, R.D. Stevens, O.R. Ilkayeva, M.J. Brosnan, T.P. Rolph, P.A. Grimsrud, D.M. Muoio, The Acetyl Group Buffering Action of Carnitine Acetyltransferase Offsets Macronutrient-Induced Lysine Acetylation of Mitochondrial Proteins, Cell Rep 14 (2016) 243–254. 10.1016/j.celrep.2015.12.030.

[72] A. Prokesch, H.J. Pelzmann, A.R. Pessentheiner, K. Huber, C.T. Madreiter-Sokolowski, A. Drougard, M. Schittmayer, D. Kolb, C. Magnes, G. Trausinger, W.F. Graier, R. Birner-Gruenberger, J.A. Pospisilik, J.G. Bogner-Strauss, N-acetylaspartate catabolism determines cytosolic acetyl-CoA levels and histone acetylation in brown adipocytes, Sci Rep 6 (2016) 1–12. 10.1038/srep23723.

[73] A.R. Pessentheiner, H.J. Pelzmann, E. Walenta, M. Schweiger, L.N. Groschner, W.F. Graier, D. Kolb, K. Uno, T. Miyazaki, A. Nitta, D. Rieder, A. Prokesch, J.G. Bogner-Strauss, NAT8L (N-acetyltransferase 8-like) accelerates lipid turnover and increases energy expenditure in brown adipocytes, Journal of Biological Chemistry 288 (2013) 36040–36051. 10.1074/jbc.M113.491324.

[74] V. Rubio, H.G. Britton, S. Grisolia, Mitochondrial carbamoyl phosphate synthetase activity in the absence of N-acetyl-L-glutamate. Mechanism of activation by this cofactor, Eur J Biochem 134 (1983) 337–343. 10.1111/j.1432-1033.1983.tb07572.x.

[75] V.T. Bortoluzzi, R.T. Ribeiro, Â.B. Zemniaçak, S. de A. Cunha, J.O. Sass, R.F. Castilho, A.U. Amaral, M. Wajner, Disturbance of mitochondrial functions caused by N-acetylglutamate and N-acetylmethionine in brain of adolescent rats: Potential relevance in aminoacylase 1 deficiency, Neurochem Int 171 (2023) 105631. 10.1016/j.neuint.2023.105631.

[76] V.T. Bortoluzzi, R.T. Ribeiro, C.V. Pinheiro, E.T. Castro, T.Q. Tavares, G. Leipnitz, J.O. Sass, R.F. Castilho, A.U. Amaral, M. Wajner, N-Acetylglutamate and N-acetylmethionine compromise mitochondrial bioenergetics homeostasis and glutamate oxidation in brain of developing rats: Potential implications for the pathogenesis of ACY1 deficiency, Biochem Biophys Res Commun 684 (2023) 149123. 10.1016/j.bbrc.2023.149123.

[77] R. Conrad, K. Narayan, Instance segmentation of mitochondria in electron microscopy images with a generalist deep learning model trained on a diverse dataset, Cell Syst 14 (2023) 58–71.e5. 10.1016/j.cels.2022.12.006.

[78] C.A. Pileggi, C.P. Hedges, S.A. Segovia, J.F. Markworth, B.R. Durainayagam, C. Gray, X.D. Zhang, M.P.G. Barnett, M.H. Vickers, A.J.R. Hickey, C.M. Reynolds, D. Cameron-Smith, Maternal high fat diet alters skeletal muscle mitochondrial catalytic activity in adult male rat offspring, Front Physiol 7 (2016). 10.3389/fphys.2016.00546.

[79] W. Guo, L. Jiang, S. Bhasin, S.M. Khan, R.H. Swerdlow, DNA extraction procedures meaningfully influence qPCR-based mtDNA copy number determination, Mitochondrion 9 (2009) 261–265. 10.1016/j.mito.2009.03.003.

[80] K.J. Livak, T.D. Schmittgen, Analysis of Relative Gene Expression Data Using Real-Time Quantitative PCR and the 2−ΔΔCT Method, Methods 25 (2001) 402–408.

[81] W. Lu, M.F. Clasquin, E. Melamud, D. Amador-Noguez, A.A. Caudy, J.D. Rabinowitz, Metabolomic analysis via reversed-phase ion-pairing liquid chromatography coupled to a stand alone orbitrap mass spectrometer, Anal Chem 82 (2010) 3212–3221. 10.1021/ac902837x.

[82] Z. Pang, Y. Lu, G. Zhou, F. Hui, L. Xu, C. Viau, A.F. Spigelman, P.E. Macdonald, D.S. Wishart, S. Li, J. Xia, MetaboAnalyst 6.0: towards a unified platform for metabolomics data processing, analysis and interpretation, Nucleic Acids Res 52 (2024) W398–W406. 10.1093/nar/gkae253.

